# Neural basis of approximate number system develops independent of visual experience

**DOI:** 10.1101/573436

**Authors:** Shipra Kanjlia, Lisa Feigenson, Marina Bedny

**Author notes:** Corresponding Author 3400 N Charles St. Ames 232 Baltimore, MD, 21218.

## Abstract

Thinking about numerical quantities is an integral part of daily human life that is supported by the intraparietal sulcus (IPS). The IPS is recruited during mathematical calculation and neuronal populations within the IPS code for the quantity of items in a set. Is the developmental basis of IPS number representations rooted in visual experience? We asked if the IPS possesses population codes for auditory quantities in sighted individuals and, critically, whether it does in the absence of any visual experience in congenitally blind individuals. We found that sequences of 4, 8, 16 and 32 tones each elicited unique patterns of fMRI activity in the IPS of both sighted and congenitally blind individuals, such that the quantity a participant heard on a given trial could be reliably predicted based on the pattern of observed IPS activity. This finding suggests that the IPS number system is resilient to dramatic changes in sensory experience.

## Introduction

Numerical reasoning is often associated with formal mathematics involving the ability to manipulate symbols to perform exact calculations. These mathematical activities constitute a uniquely human domain of knowledge that has developed recently in human history. However, aspects of mathematical reasoning are thought to build upon a more evolutionarily ancient foundation of non-symbolic number reasoning that is shared with other species (Dehaene & Cohen, 2007; Dehaene, Piazza, Pinel, & Cohen, 2003; Feigenson, Dehaene, & Spelke, 2004), and that is present in the first days of human life (Izard et al., 2009). This intuitive number sense enables humans and non-human animals to estimate the approximate quantity of items in sets, such as the approximate number of people in a room, or the approximate number of fruits in a foraging patch.

A key feature of the Approximate Number System (ANS) is that its representations of quantity are noisy. Without language to enable exact counting, it is not possible to determine whether there are exactly 16 or 17 people in a room. Instead, a given magnitude is represented by a distribution of neural activity that is centered upon the true magnitude but is also characterized by some amount of variance. Thus, a set of 16 will often be mistaken for 15 or 17 items. The variance in approximate number representations has been hypothesized to increase linearly with distribution means—a feature termed scalar variability. This property predicts that a pair of smaller magnitudes, such as 8 and 12, will be more discriminable than a pair of larger magnitudes that are equally numerically distant, such as 30 and 34, because smaller magnitudes are characterized by less variability. Since variability scales with numerosity, the discriminability of two quantities depends specifically upon their ratio and not their absolute magnitude difference (e.g. 8 vs. 12 and 30 vs. 45). This feature gives rise to the approximate number system’s key ratio-dependent behavioral signature (Feigenson et al., 2004).

In humans, both symbolic and non-symbolic approximate numerical processing are supported by fronto-parietal circuits (Holloway, Price, & Ansari, 2010; Piazza & Eger, 2016; Piazza, Mechelli, Price, & Butterworth, 2006; Piazza, Pinel, Le Bihan, & Dehaene, 2007; Venkatraman, Ansari, & Chee, 2005). The intraparietal sulcus (IPS) in particular seems to play a crucial role in numerical processing. An especially compelling piece of evidence for this idea comes from findings of neuronal populations codes for specific quantities in the IPS of monkeys and humans. Intracranial recordings from the IPS of monkeys have uncovered “number neurons” that are tuned to specific quantities, even in the absence of training (Nieder, 2005, 2012; Nieder, Diester, & Tudusciuc, 2006; Nieder, Freedman, & Miller, 2002; Viswanathan & Nieder, 2013). In humans, repeated presentations of visual sets of the same quantity causes neural adaptation in the IPS with ratio-dependent recovery in activity following deviant quantities (Piazza, Pinel, Bihan, Dehaene, & Cedex, 2004; Piazza et al., 2007). Furthermore, numerical magnitudes elicit different spatial patterns of activity in the IPS (Bluthe, De Smedt, & Op de Beeck, 2015; Cavdaroglu, Katz, & Knops, 2015; Eger et al., 2009; Harvey, Klein, Petridou, & Dumoulin, 2013; Harvey & Dumoulin, 2017; Lyons, Ansari, & Beilock, 2014). For example, sets of 4, 8, 16 or 32 objects evoke different patterns of IPS activity. After training, a linear support vector machine can use these IPS activity patterns to predict which quantity the participant viewed on a given trial (Eger et al., 2009). Furthermore, consistent with behavioral approximate number discrimination, IPS activity patterns for quantities demonstrate ratio-dependency--quantities that differ by a smaller ratio (e.g. 4 and 8 vs. 4 and 16) evoke more overlapping neural patterns in the IPS (Eger et al., 2009). Together these data support the idea that neuronal populations in the IPS code for numerical magnitude.

A key open question concerns the developmental origins and role of experience in the establishment of IPS quantity representations. IPS responses to approximate number are present in children by the age of 3 to 4 years (Cantlon, Brannon, Carter, & Pelphrey, 2006; Kersey & Cantlon, 2017), and parietal responses to numerosities are revealed by near-infrared spectroscopy in 6-month old infants (Hyde, Boas, Blair, & Carey, 2010) and event-related potentials in 3-month old infants (Izard, Dehaene-Lambertz, & Dehaene, 2008). However, it is unclear whether and what kind of experience is necessary for this specialization.

One possibility is that IPS representations of number develop as a result of accumulated experience with visual sets. There is some evidence that vision has a privileged status for conveying numerical information. Like early visual features, such as color, contrast and orientation, number induces after-effects. Prolonged exposure to a large number of dots causes participants to underestimate a subsequent set of dots (Burr & Ross, 2008; Ross, 2010), suggesting that the visual system is capable of sensing number directly (Burr & Ross, 2008; Ross, 2010). Furthermore, neural networks trained with images of arrays containing different numbers of objects spontaneously develop representations of numerosity (Stoianov & Zorzi, 2012). That is, “neurons” in the hidden layer of the model develop selectivity to number, akin to the number-neurons found in monkey IPS (Stoianov & Zorzi, 2012). IPS neurons may similarly develop responses to quantity through experience with visual arrays.

On the other hand, there are reasons to think that numerical representations in the IPS are abstract, rather than visual. First, numerical information can be represented from non-visual input; adults can approximate the number of events in auditory sequences and enumerate points of tactile stimulation (Gallace, Tan, & Spence, 2006, 2008; Piazza et al., 2006; Riggs et al., 2006; Tokita, Ashitani, & Ishiguchi, 2013). Second, from infancy onward there is integration between visual and non-visual numerical information. Newborn infants can numerically match the numbers of visual and auditory stimuli (Izard, Sann, Spelke, & Streri, 2009), and 6-month old infants can use auditory information to predict the number of visual items in an array (Feigenson, 2011). Both adults and children can compare approximate quantities across modalities (e.g., judging whether there are more items in a visual array or in an auditory sequence) (Barth, Kanwisher, & Spelke, 2003; Barth, La Mont, Lipton, & Spelke, 2005). Finally, both visual and auditory numerical stimuli, whether non-symbolic or symbolic, all activate the IPS (Eger, Sterzer, Russ, Giraud, & Kleinschmidt, 2003; Piazza et al., 2006). Nevertheless, it remains possible that vision “bootstraps” representations of quantity in the IPS. Once in place, IPS number representations could become accessible through other sensory modalities and, in numerate humans, through symbols such as Arabic numerals. Neuroimaging findings in children do not address this possibility, as they have focused on visual stimuli (Cantlon et al., 2006; Izard et al., 2008; Kersey & Cantlon, 2017).

It is now known that vision is not required for the development of symbolic number responses in the IPS. In both congenitally blind and sighted individuals, the IPS is more active during math calculation than sentence comprehension and demonstrates sensitivity to the number of digits and algebraic complexity of math equations (Amalric, Denghien, & Dehaene, 2017; Crollen et al., 2019; Kanjlia, Lane, Feigenson, & Bedny, 2016). However, these findings leave open the question of whether the neural basis of our basic approximate number system is preserved in the absence of vision. Prior evidence suggests that, within the IPS, responses to approximate and exact number are neurally separable (Bluthe et al., 2015; Damarla & Just, 2013; Dehaene, Spelke, Pinel, Stanescu, & Tsivkin, 1999; Lyons et al., 2015). Machine-learning classifiers that are trained on patterns of activity evoked by symbolic quantities are not able to discriminate between patterns evoked by non-symbolic quantities (Bluthe et al., 2015; Damarla & Just, 2013; Lyons et al., 2015). Therefore, whether IPS responses to approximate number develop in the total absence of vision remains an open question. Furthermore, even if the IPS develops representations of approximate number without visual experience, it is possible that the precision of these representations is refined by visual experience. If the precision of approximate number representations is honed by visual experience, quantity representations may be less discriminable in the IPS of congenitally blind individuals relative to sighted individuals.

Here we asked whether visual experience is necessary for the development of approximate number representations in the IPS. Our key question is whether the IPS of congenitally blind individuals is sensitive to the number of events in auditory sequences and whether these quantities are coded in the IPS in a ratio-dependent manner. We trained a machine-learning classifier on IPS activity patterns associated with listening to sequences of 4, 8, 16 or 32 tones, and asked whether it could predict the number of tones a participant heard on a given trial. Furthermore, we asked whether the IPS represents quantities with similar precision in both congenitally blind and sighted individuals. We tested whether quantities could be decoded from IPS activity with similar accuracy across congenitally blind and sighted individuals, and whether ratio had a similar effect on classification accuracy across the two groups. If visual experience is necessary for improving the precision of IPS quantity representations, discrimination accuracy of the classifier should be lower in the congenitally blind compared to the sighted group.

A second goal of the current study was to investigate potential cross-modal recruitment of the visual cortex for approximate number processing. In addition to the IPS, congenitally blind but not sighted individuals recruit parts of dorsal occipital cortex, specifically the right middle occipital gyrus (rMOG), during math calculation (Amalric et al., 2017; Crollen et al., 2019; Kanjlia et al., 2016). Furthermore, like the IPS, the math-responsive area of the “visual” cortex in blind participants was sensitive to the number of digits and algebraic complexity of math equations (Kanjlia et al., 2016). One hypothesis is that parts of the “visual” cortex of congenitally blind individuals are repurposed for higher-cognitive functions such as numerical processing, potentially via top-down connectivity with fronto-parietal networks (Bedny, 2017). Indeed, even in the absence of a task (i.e. during rest), math-responsive “visual” cortex, rMOG, shows increased synchrony with the fronto-parietal number network in congenital blindness (Kanjlia et al., 2016). Therefore, if the rMOG is incorporated into the fronto-parietal number network in congenital blindness, it should also demonstrate sensitivity to numerical approximation. Here we asked whether math-responsive rMOG codes for approximate numerical quantities in a manner similar to the IPS.

## Materials & Methods

### Participants

Seventeen congenitally blind (12 female, mean age 49 years, SD=16, min=29, max=73) and twenty-five sighted control participants (16 female, mean age 43 years, SD=16, min=19, max=67) contributed data to the final sample. Five additional participants were tested but excluded from the final sample because further screening revealed a history of some vision (1 blind participant), because they fell asleep during the experiment (2 sighted participants) or due to poor performance (2 sighted participants). Blind participants had at most minimal light perception and their blindness was due to pathology of the eyes or optic nerve, not due to brain damage. All participants reported having no cognitive or neurological disorders.

**Table 1.** Participant demographic information.

| **Subject Number** | **Gender** | **Age** | **Cause of Blindness** | **Light Perception** | **Education** |
| --- | --- | --- | --- | --- | --- |
| 1 | M | 50 | LCA | None | JD |
| 2 | F | 35 | ROP | Minimal | BA |
| 3 | F | 38 | LCA | Minimal | MA |
| 4 | M | 31 | Anophthalmia | None | SC |
| 5 | F | 42 | ROP | None | MA |
| 6 | F | 67 | ROP | None | MA |
| 7 | M | 47 | Unknown | Minimal | BA |
| 8 | F | 29 | ROP | Minimal | MA |
| 9 | F | 64 | ROP | None | HS |
| 10 | F | 73 | ROP | Minimal | HS |
| 11 | M | 46 | Unknown | None | JD |
| 12 | M | 70 | ROP | Minimal | HS |
| 13 | F | 31 | Detached Optic Nerve | None | SC |
| 14 | F | 65 | RP | Minimal | BA |
| 15 | F | 29 | LCA | Minimal | BA |
| 16 | F | 64 | ROP | None | JD |
| 17 | F | 55 | Cataracts, Microphthalmia | None | PhD |
| **Average** |  |  |  |  |  |
| **Blind**<br>**(n=17)** | **12 F** | **49**<br>**(SD=16)** | **--** | **--** | **BA** |
| **Sighted**<br>**(n=25)** | **16 F** | **43**<br>**(SD=16)** | **--** | **--** | **BA** |

LCA=Leber Congenital Amaurosis; ROP=Retinopathy of Prematurity; RP=Retinitis Pigmentosa; BA=Bachelor of Arts; HS=High School; JD=Juris Doctor; MA=Master of Arts; SC=Some College

### fMRI Task

Participants completed an auditory approximate number comparison task that was adapted from a visual approximate number comparison task designed by Eger et al. (2009). On each trial, participants first heard a tap to indicate the trial was starting. This was followed by a sample sequence of 4, 8, 16 or 32 beeps. After a 6-second delay, they heard a second, test sequence of beeps whose numerosity differed from the first sequence by a ratio of 2 (e.g. sample sequence: 8 beeps, test sequence: 4 or 16 beeps). The test sequence never exceeded 32 beeps and was never smaller than 4 beeps. After a second tap (to indicate the end of the second stimulus), participants had 4 seconds to indicate whether the second sequence was more or less numerous than the first by pressing one of two buttons. Each trial was followed by a 6-second rest period. Participants were instructed to try not to count the beeps but rather estimate the number of beeps.

Only neural activity during the first, sample sequence was used for further analyses. To ensure that numerosity was not being coded on the basis of low-level stimulus features, sample sequences were matched across numerosities either on total sequence duration or individual element duration (i.e. 2 match conditions). In the total duration matched condition, the sample sequence for every numerosity was 3 seconds long, with larger numerosities (e.g. 32) playing faster than smaller numerosities (e.g. 4). In the element duration matched condition, each beep in the sample sequence played for ∼0.2 seconds (with some jitter), thus matching numerosities on pace but not overall duration.

To discourage participants from using the duration of the sequences as a cue to numerosity, the duration of the test sequences was either congruent or incongruent with respect to the numerical ratio between the sample and test sequence. On congruent trials, test sequences that were more numerous than sample sequences played twice as long, and those that were less numerous were half as long (e.g., sample sequence: 8 beeps, 3 seconds; test sequence: 16 beeps, 6 seconds), and vice versa on incongruent trials. Note, however, that neural responses to the second, test sequences were not analyzed.

Participants were not informed about the range of numerosities, the match conditions or the manipulation of congruence. However, they were told that the speed and duration of the beeps may vary and were instructed to ignore these features and attend to the quantity of beeps. Rather than trial-by-trial feedback, participants were given a score (percent correct) at the end of each run of the task.

The entire experiment was broken up into 8 sections, or runs. Each of the 8 sample conditions (4 numerosities by 2 match conditions) appeared on 4 trials per run (32 total trials per run). The 8 sample conditions were arranged in a Latin Square design such that each condition followed and preceded every other condition an equal number of times over the course of the experiment.

Both blind and sighted participants were blind-folded throughout. All participants completed all 8 runs of the experiment.

### Data Acquisition and Univariate Analysis

MRI data were collected using a 3T Phillips scanner. Structural images were T1-weighted and were collected in 150 axial slices (1-mm isotropic voxels). Functional data sensitive to BOLD contrast were collected in 36 axial slices (2.4 × 2.4 × 3 mm voxels; repetition time 2 seconds). MRI data were analyzed using Freesurfer, FSL, HCP workbench and custom in-house software.

Preprocessing steps included motion correction, high-pass filtering (128 s), mapping the data to the cortical surface using Freesurfer, smoothing with a 6mm FWHM Gaussian kernel on the surface, and prewhitening to remove temporal autocorrelation.

Data were then analyzed using a general linear model, which included eight regressors of interest—one for each sample condition (4 numerosities by 2 match conditions) that modeled the first stimulus and delay periods together. Resulting z-statistic maps for each of the 8 regressors of interest for each run were used for MVPA. The test sequence, response period as well as the instruction taps (prior to first stimulus and prior to second stimulus) were modeled separately and were not included in any of the reported analyses. We also separately modeled and excluded trials in which the participant failed to respond.

### Multi-voxel Pattern Analysis (MVPA)

We used MVPA to ask whether the following four regions of interest (ROIs) contained a spatial code for auditory numerosities: right IPS, left IPS, right middle occipital gyrus (rMOG) within visual cortex, and early auditory cortex (A1). Group-specific IPS ROIs were defined based on a math equations>sentences contrast from a separate published dataset (p<0.01, uncorrected for sighted and p<0.001, uncorrected for congenitally blind) (see Kanjlia et al., 2016 for details). Briefly, in that experiment, participants heard pairs of math equations each with a variable x, and had to judge whether x had the same value in two equations. In the control condition, they judged whether a pair of sentences, one in the passive voice and one in the active voice, had the same meaning. The math>sentences contrast in this experiment identified bilateral math-responsive IPS ROIs in both sighted and blind individuals. Additionally, in the blind group only, responses to math were observed in the rMOG of the “visual” cortex. These ROIs were used in the current study.

The math-responsive “visual” cortex ROI (rMOG) was defined as the cluster within the right visual cortex that responded more to math equations than sentences in congenitally blind>sighted individuals (right middle occipital gyrus, rMOG; p<0.01, uncorrected) (Kanjlia et al., 2016). To ask whether the auditory cortex was sensitive to numerosity, we used a previously published auditory cortex ROI that includes anatomically defined posteromedial, middle and anterolateral Heschel’s gyrus (Norman-Haignere et al. 2013).

MVPA was conducted using the pyMVPA toolbox (Hanke et al., 2009). We used MVPA to decode numerosity (6 total comparisons: 4 vs. 8, 4 vs. 16, 4 vs. 32, 8 vs. 16, 8 vs. 32, 16 vs. 32) based on patterns of activity within each ROI using a leave-one-run-out cross-validation procedure. For a given pair of numerosities (e.g., 4 vs. 8), a linear support vector machine (SVM) was trained on 28 vectors of neural activity (2 numerosities x 2 match conditions x 7 runs) within an ROI and then tested on 4 vectors (2 numerosities x 2 match conditions) from the left-out run. This process was repeated iteratively until every run was left out and classification accuracy was averaged over cross-validation folds. To evaluate overall classification performance, we averaged classification accuracy over all 6 numerosity pairs.

We further asked whether regions that code for numerosity demonstrate a known signature of the approximate number system: ratio-dependent numerosity coding. Quantities that differ by a smaller ratio are known to be harder to distinguish behaviorally and also activate more overlapping neuronal populations (Feigenson et al., 2004; Nieder, 2013; Piazza et al., 2004, 2007; Viswanathan & Nieder, 2013). Thus, we predicted that regions that code for quantity would demonstrate more closely overlapping neural patterns (i.e., lower classification accuracy) for quantities that differ by smaller ratios than larger ratios. We compared classification performance across pairs of different ratios, collapsing over pairs of numbers that differed by the same ratio (e.g. 4 vs. 8 and 8 vs. 16 are both ratio 2). Ratio effects were statistically tested using a repeated measures ANOVA with ratio as a covariate and hemisphere and group as categorical factors.

Next we compared the degree to which numerosity decoding was driven by numerical as opposed to non-numerical, low-level stimulus features across ROIs. Numerosity was more confounded with overall amount of sound on trials that were matched on individual beep duration than those matched on overall duration. Therefore, we compared classification accuracy for element-duration and total-duration matched quantities separately, in order to test the hypothesis that the auditory cortex (A1) is more sensitive to total amount of sound rather than actual numerosity, whereas the IPS is sensitive to numerosity, even when overall amount of sound is controlled. The same test was also conducted within the “visual” rMOG ROI.

Finally, we used a searchlight analysis to ask where quantities could be decoded across the entire cortex. For each participant and pair of numerosities, MVPA was conducted within searchlight regions of 10mm radius across the cortical surface. Classification accuracy across all 6 quantity pairs was then averaged within each searchlight. Accuracy maps were logit-transformed and then statistically compared across participants within a group and across groups using random-effects GLM analyses.

## Results

### Behavioral Results

Behaviorally, both sighted and congenitally blind groups performed well above chance and no differently from each other (sighted: 85.28%, SD=9.02%; congenitally blind: 88.24%, SD=1.10; t(41)=0.58, p=0.57).

### Similar ratio dependent sensitivity in IPS of sighted and blind

Within the left and right math-responsive IPS of the sighted group, the classifier discriminated patterns evoked by the presented auditory quantities (i.e. 4, 8, 16 and 32) above chance (left 56.04% (SD=1.48), one-sample t-test t(24)=4.12, p<0.001; right IPS 57.29% (SD=1.44) one-sample t-test t(24)=5.26, p<0.001; paired t-test between hemispheres: t(24)=-1.21, p=0.24) (Fig. 1). Furthermore, numerosities that differed by a larger ratio were discriminated with higher accuracy (main effect of ratio: F(1,122)=27.04, p<0.001; main effect of hemisphere: F(1,122)=1.59, p=0.21; ratio by hemisphere interaction: F(1,122)=0.74, p=0.39).

**Fig. 1.**
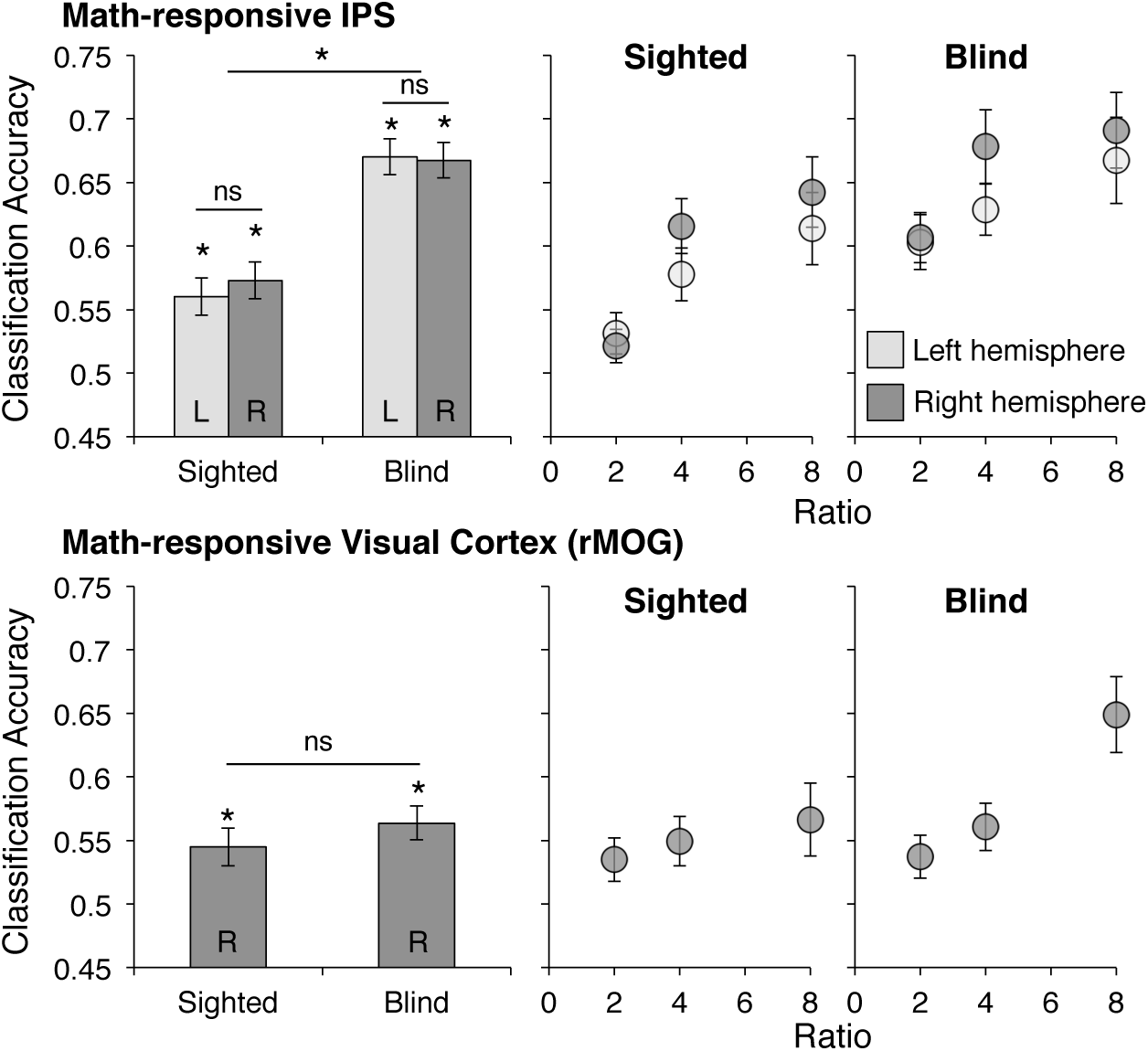
MVPA Classification accuracy in math-responsive IPS and rMOG. Linear SVM accuracy for classifying neural patterns in the left and right IPS (top panel) and rMOG (bottom panel). Classification accuracy is averaged across all numerosity pairs in bar graphs and is averaged across all numerosity pairs of the same ratio in scatter plots.

Similarly, in the congenitally blind group, the classifier discriminated activity patterns in left and right math-responsive IPS with 61.82% (SD=2.01) and 64.29% (SD=1.80) accuracy, respectively (left: t(16)=6.33, p<0.001; right: t(16)=8.51, p<0.001; paired t-test between hemispheres: t(24)=-1.48, p=0.16) (Fig. 1). Analogous to the sighted group, there was an effect of numerical ratio on decoding accuracy (main effect of ratio: F(1,82)=10.54, p=0.002; main effect of hemisphere: F(1,82)=2.10, p=0.15, ratio by hemisphere interaction: F(1,82)=0.08, p=0.78).

Overall decoding accuracy was better in the IPS of congenitally blind than sighted individuals (hemisphere by group repeated measures ANOVA; main effect of group: F(1,40)=9.99, p=0.003). The effect of ratio did not differ across groups (main effect of ratio: F(1,204)=37.05, p<0.001; ratio by group interaction: F(1,204)=0.77, p=0.38; main effect of hemisphere: F(1,204)=3.57, p=0.06).

### Math-responsive visual cortex (rMOG) shows effect of ratio on decoding accuracy in congenitally bind group

In previous work we found that regions in the right “visual” cortex (right middle occipital gyrus, rMOG) were recruited during math calculation in congenitally blind but not sighted individuals (Kanjlia et al., 2016). Here we found that non-symbolic auditory numerosities can be decoded in the rMOG of both blind and sighted participants (blind rMOG quantity decoding 56.37% (SD=1.32, one-sample t-test; t(16)=4.83, p<0.001; sighted 54.50%, SD=1.49, t(24)=3.16, p=0.004) (Fig. 1). Although decoding accuracy was slightly better in the blind group, the between-group difference in overall classification accuracy was not significant (CB vs. S: t(40)=0.92, p=0.30). However, the classifier only showed the ratio-dependent signature of quantity discrimination in the rMOG of congenitally blind not sighted individuals (main effect of ratio in CB: F(1,33)=15.83, p=0.004; S: F(1,49)=1.29, p=0.26, Fig. 1). Direct comparison of congenitally blind and sighted individuals revealed that the effect of ratio was significantly greater in the congenitally blind than the sighted group (ratio by group interaction: F(1,82)=4.37, p<0.05; main effect of group: F(1,40)=2.03, p=0.16).

### Greater effect of low-level auditory features on decoding within A1 than IPS or “visual” rMOG

Apart from the IPS and rMOG, auditory quantities were also discriminable in the auditory cortex (A1) of congenitally blind (66.90%, SD=1.11) and sighted adults (67.03% (SD=1.15) (between group t-test: t(40)=-0.83, p=0.93).

To test whether decoding was driven by low-level features (i.e., overall amount of sound) to a greater extent in auditory cortex (A1) than in the IPS or rMOG, we compared decoding performance across element-matched and duration-matched conditions. In the element matched condition, quantities with greater numerical distances also differed from each other in overall amount of sound; this was not the case in the duration-matched sequences. We therefore reasoned that cortical areas that were more sensitive to overall amount of sound than numerical quantity would show better decoding performance for the element matched than the duration matched condition. Consistent with the idea that decoding in A1 was driven more by overall amount of sound, the difference in decoding accuracy between element-matched and duration matched lists was more pronounced in A1 than in the IPS in both the sighted and blind groups (hemisphere by match-condition by ROI repeated-measures ANOVA; ROI by match-condition interaction in sighted group: F(1,24)=69.25, p<0.001; blind group: F(1,16)=15.62, p=0.001) (Fig. 2). Analogously, in the both groups, the difference between element-matched and duration matched lists was more pronounced in right A1 than rMOG (match-condition by ROI repeated-measures ANOVA; main effect of match-condition in sighted group: F(1,24)=56.47, p<0.001; blind group: F(1,16)=11.31, p=0.004).

**Fig. 2.**
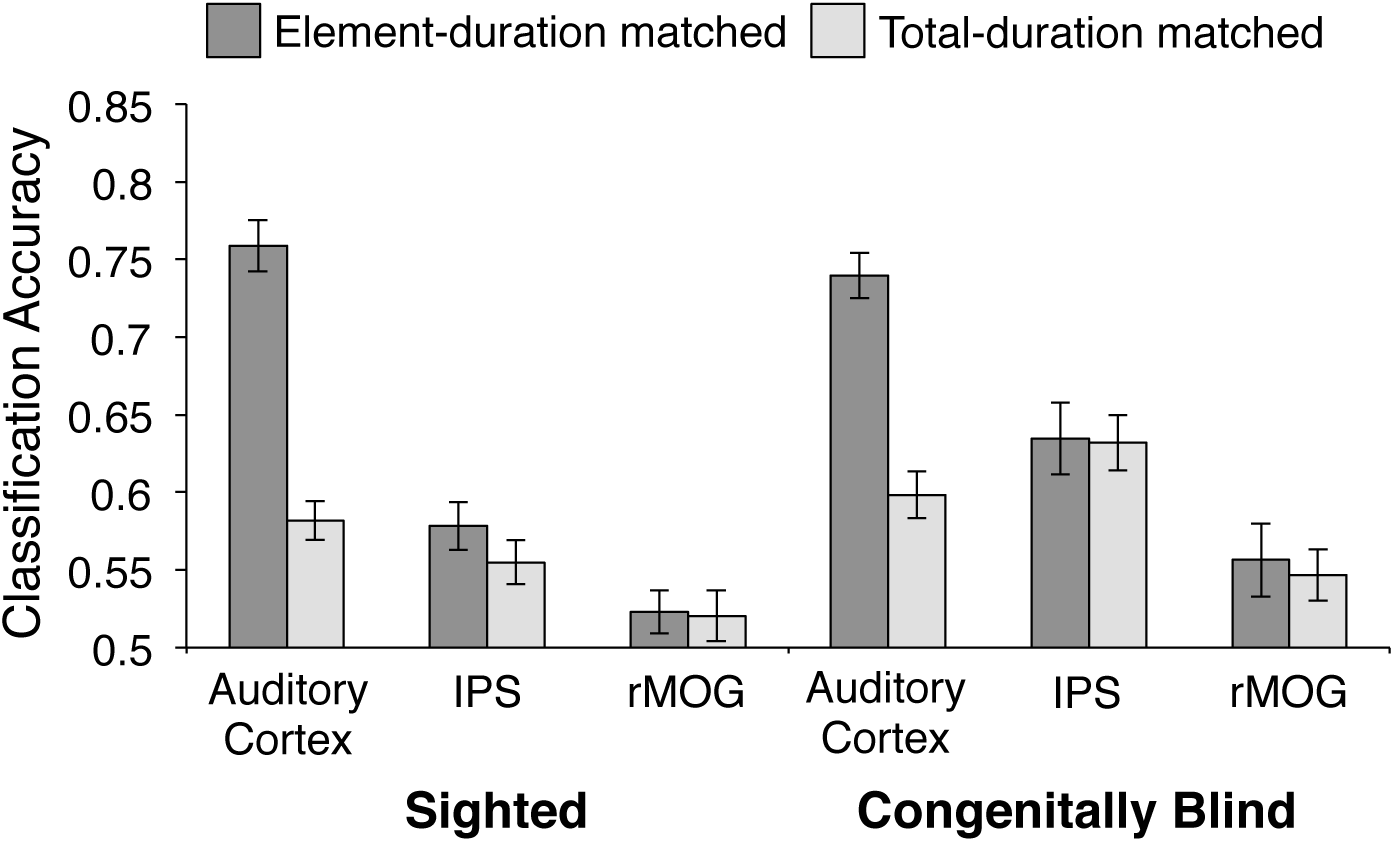
Effect of low-level auditory features on MVPA classification accuracy. Linear SVM discrimination performance for stimuli that were matched on either element duration or total duration across auditory cortex, IPS and rMOG ROIs. Classification accuracy is averaged across left and right hemispheres for auditory cortex and IPS ROIs.

### Searchlight analyses reveal auditory quantity decoding in fronto-parietal number network

Searchlight analyses revealed successful decoding of auditory quantities in a right-lateral fronto-parietal network in both congenitally blind individuals and sighted (Fig. 3). In congenitally blind individuals, numerosity decoding extended posteriorly along the dorsal occipital cortex (rMOG) as well as lateral occipito-temporal cortex, in the vicinity of the visual number form area (VNFA). However, direct comparison of searchlight results across congenitally blind and sighted groups only yielded significant differences in the right superior frontal sulcus and gyrus (Fig. 3; Supplementary Table 1).

**Fig. 3.**
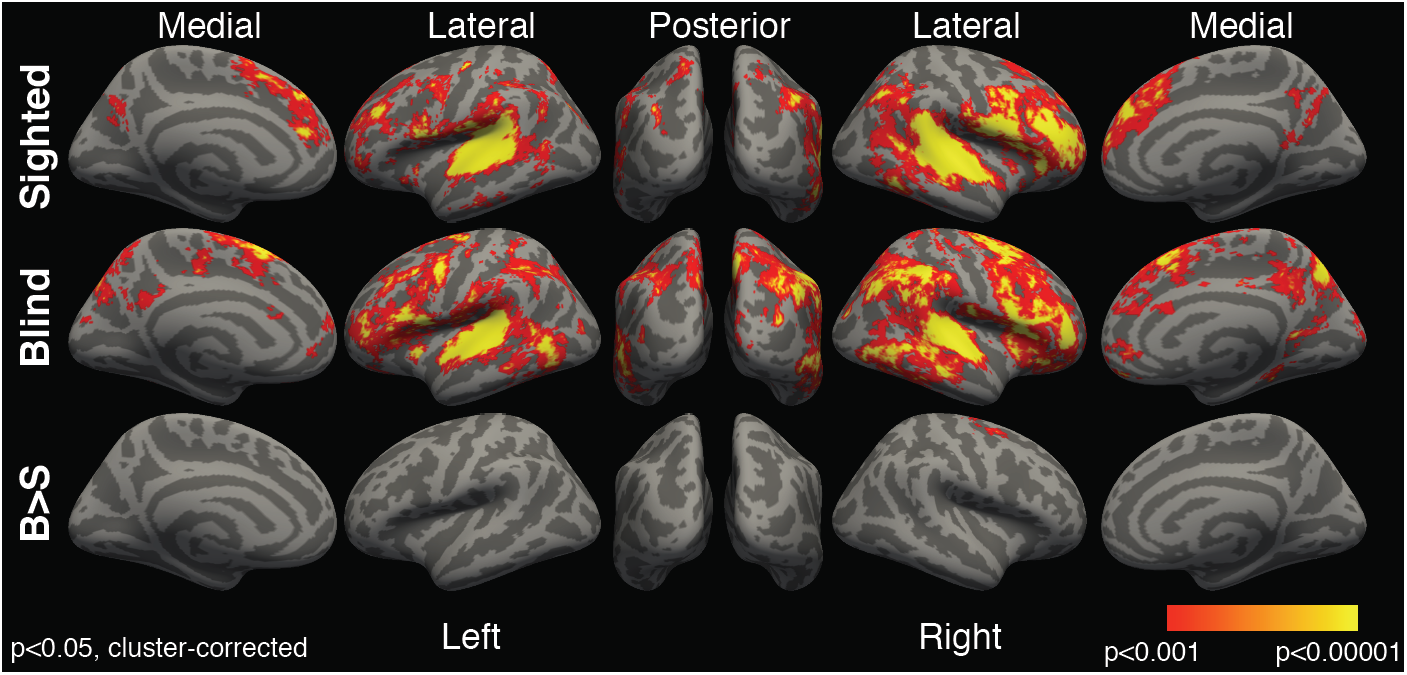
Numerosity classification whole-cortex searchlight results, cluster corrected p<0.05, vertex-wise threshold p<0.001.

## Discussion

### Representations of number in the IPS are modality independent

Previous studies with sighted individuals have found that spatial patterns of activity within the IPS discriminate between different numerical quantities. That is, when sighted participants view arrays of visual objects (e.g., dots), the pattern of activity within the IPS reflects the approximate number of items seen (Eger et al., 2009). Here we report that the IPS of sighted individuals also codes for auditory quantities. When sighted participants listened to sequences of tones, the numerosity of the tones could be decoded from spatial patterns of activity within the IPS. This finding replicates the results of one previous fMRI study, which found that activity patterns in the IPS can be used to discriminate between auditory sequences of different quantities (Cavdaroglu et al., 2015).

As in previous work, we found that numerosity could also be decoded from activity patterns in primary auditory cortex, A1 (Cavdaroglu et al., 2015). However, relative to the IPS, decoding in A1 was more influenced by low-level properties of the stimuli (i.e., overall amount of sound). The present results from sighted participants go one step beyond previous findings by showing that the coding of auditory numerosities in the IPS is ratio-dependent. We find that quantities that are more similar to each other (differ by a smaller ratio) elicited more overlapping neural patterns in the IPS than quantities that differ by a larger ratio. This ratio-dependence of the IPS population code mirrors the ratio-dependence of behavioral discrimination and neural responses to number reported in previous work (Eger et al., 2009; Odic, Libertus, Feigenson, & Halberda, 2013; Piazza et al., 2007; Tokita et al., 2013).

The present results thus suggest that the IPS contains modality-independent representations of number. Consistent with this idea, electrophysiological recordings from the IPS of monkeys find overlap between auditory and visual representations of number at the level of individual neurons (Nieder, 2012). That is, some neurons in the IPS are tuned to a specific number of events in a sequence, regardless of whether the events are experienced through vision or audition (Nieder, 2012). However, there is also evidence that the IPS additionally has specialized representations for auditory versus visual numerical information. Single-unit recordings from neurons in the IPS show that, whereas some neurons are tuned to the same numerical magnitude across presentation formats, the majority are modality specific (Nieder, 2012). Thus, the available evidence suggests that humans too develop both modality-independent and modality-specific representations of number, but all of these representations share a neuroanatomically similar location in the IPS.

### IPS representations of number and visual experience

A key finding of the current study is that all of the functional signatures of IPS number responses that have been identified in sighted participants are also present in people who are blind from birth. The IPS of congenitally blind individuals shows ratio-dependent numerosity coding that is less sensitive to low-level auditory features than A1. This finding is consistent with behavioral studies that show preserved signatures of numerical reasoning in congenital blindness. Congenitally blind individuals show similar or slightly better performance than sighted individuals when estimating numbers of tones, footsteps or finger taps, and performance is ratio-dependent in both groups (Castronovo & Delvenne, 2013; Castronovo & Seron, 2007; Kanjlia, Feigenson, & Bedny, 2018). Prior studies also find that, like people who are sighted, individuals who are congenitally blind recruit the IPS during symbolic number reasoning and show similar behavioral correlations between numerical approximation and symbolic math performance across individuals (Amalric et al., 2017; Crollen et al., 2019; Kanjlia, Feigenson, et al., 2018; Kanjlia et al., 2016). Together, these findings suggest that numerical representations are established in the IPS independent of gross differences in sensory experience.

Evolutionary precursors for numerical processing could make the IPS robust to large-scale sensory change. The ability to approximate number is present in humans from birth and is shared with various species, including non-human primates, rats, birds and fish (Agrillo, Dadda, Serena, & Bisazza, 2008; Cantlon & Brannon, 2006; Izard et al., 2009; Meck & Church, 1983; Roberts, Coughlin, & Roberts, 2000). Homologous areas of the brain support numerical processing in humans and non-human primates (Nieder, 2013; Viswanathan & Nieder, 2013). These findings suggest that the seeds of numerical reasoning are present in our evolutionary heritage.

Although the basic signatures of IPS selectivity are similar in congenital blindness, the present results also provide some evidence for enhanced responses to auditory number sequences in the IPS and prefrontal cortices of blind individuals. Numerosities were decoded with 12% more accuracy in the IPS of congenitally blind relative to sighted individuals. In whole-cortex analysis, we also observed a greater extent of auditory quantity decoding across the cortex of the blind than the sighted group, particularly within the prefrontal cortex. Furthermore, previous behavioral studies find superior performance on some approximate number tasks among blind individuals, such as on production tasks and estimation of large numerosities (Castronovo & Delvenne, 2013; Castronovo & Seron, 2007).

One possibility is that the IPS becomes more tuned to sequential auditory quantities in individuals who are blind. As noted above, while some number-responsive neurons in the IPS of monkeys are modality-and format-independent, a subset are modality-and format-specific (Nieder, 2012; Nieder et al., 2006). For example, some IPS number neurons prefer quantities presented auditorily while others prefer visual quantities (Nieder, 2012). Similarly, some number neurons respond preferentially to numerosities presented sequentially as opposed to simultaneously (e.g., 4 dots presented in sequence vs. concurrently) (Nieder et al., 2006). Blindness may enhance the proportion of number neurons in the IPS preferring auditory input, or sequentially presented input, or may selectively improve the tuning of these neurons. Some evidence suggests that the precision of approximate number representations is refined over the first few months of life in sighted individuals (Libertus & Brannon, 2010). It is possible that this tuning is modality-specific. Consistent with this idea, previous studies find that some aspects of auditory processing are enhanced in congenital blindness, such as the ability to localize sounds in space (Collignon et al., 2011). The ability to extract numerical information from auditory sequences may be similarly enhanced in congenital blindness.

According to this hypothesis, experience builds on innately specified numerical representations, enhancing their precision and modifying them to efficiently extract numerical information from the types of stimuli that are most often encountered. As noted above, infants have the ability to estimate quantities just hours after birth (Izard et al., 2009). Thus, it appears that experience is not necessary to establish representations of number, but may influence the tuning of number representations. For example, the acquisition of language and education are experiences that are known to change representations of number. Acquiring an exact, symbolic number system enables humans to encode and remember the cardinality of a set, manipulate it in the absence of a physical reference, and perform mathematical operations over it (Frank, Fedorenko, & Gibson, 2008; Spaepen, Coppola, Spelke, Carey, & Goldin-Meadow, 2011). Furthermore, acquisition of number words and mathematical education improves precision on approximate number tasks (Piazza, Pica, Izard, Spelke, & Dehaene, 2013; Pica, Lemer, Izard, & Dehaene, 2004). Humans who possess a limited vocabulary for numbers show lower precision on approximate number tasks, and this precision improves when number words are acquired (Piazza et al., 2013; Pica et al., 2004). Taken together, these findings suggest that representations of number are present from birth but can be modified by experience.

### Math-responsive visual cortices code for non-symbolic quantities in congenital blindness

We find that, in addition to the IPS, parts of the “visual” cortex, in particular the right middle occipital gyrus (rMOG), shows ratio-dependent coding of numerosity in blind but not sighted individuals. Furthermore, like the IPS, the rMOG was less sensitive to low-level auditory features than early auditory cortex (A1).

The present findings are consistent with prior evidence that the rMOG acquires responses to symbolic number in blindness (Amalric et al., 2017; Crollen et al., 2019; Kanjlia et al., 2016). Like the IPS, the rMOG of blind individuals responds preferentially during math calculation than sentence comprehension and activity increases with the difficulty of math equations (Kanjlia et al., 2016). Furthermore, even during rest, activity in the rMOG is more synchronized with IPS activity in congenitally blind compared to sighted individuals. Together with the present evidence, these findings suggest that IPS representations of number expand into deaffrented visual cortices in congenital blindness. Just as the IPS responds to both symbolic and non-symbolic numerical information, the rMOG shows sensitivity to both symbolic and non-symbolic number in blindness.

Prior studies suggest that, in sighted individuals, this rMOG region is retinotopically organized and performs mid-level visual functions such as motion and object processing (Kolster, Peeters, & Orban, 2010; Larsson & Heeger, 2006; Tootell et al., 1997; Van Essen, Glasser, Dierker, Harwell, & Coalson, 2012). Furthermore, in blindfolded sighted individuals, this region does not show an above baseline response to auditory numerical stimuli (Kanjlia et al., 2016). Interestingly, however, in the current study, we found that numerosity could be decoded from activity in the rMOG of sighted individuals, although the effect was not ratio-dependent. These results are consistent with the idea that plasticity in blindness builds upon pre-existing connectivity patterns that are common to the sighted and blind. According to this hypothesis, the visual cortex has pre-existing regional biases in functional connectivity with fronto-parietal circuits in sighted and blind individuals alike (Bedny, 2017). In sighted individuals, top-down input from fronto-parietal networks to the visual cortex is outweighed by bottom-up visual input. However, in congenital blindness, these inputs have an opportunity to repurpose the visual regions with which they communicate.

Consistent with the current data, relative to other ventral visual areas, the rMOG of sighted individuals shows higher resting-state synchrony with math-responsive IPS (Kanjlia, Pant, & Bedny, 2018). It is possible that this communication enables some number-related responses in the rMOG of sighted individuals, such as the numerosity-specific patterns observed in the current study. However, when bottom up visual input is completely removed, these network-specific fronto-occipital connectivity biases are enhanced (Bedny, Pascual-Leone, Dodell-Feder, Fedorenko, & Saxe, 2011; Crollen et al., 2019; Kanjlia et al., 2016; Kanjlia, Pant, et al., 2018). Thus, despite a common pre-existing “blue-print,” early experience alters the functional properties of cortex, engendering ratio-dependent number coding in parts of cortex that do not typically represent this information. In this regard, the results are consistent with accumulating evidence that, in blindness, “visual” cortices are colonized by top-down projections from higher-cognitive networks, such as the IPS (Bedny, 2017).

What is the relationship between IPS and rMOG representations of number? One possibility is that, in congenital blindness, numerical processing becomes distributed over two regions rather than isolated to the IPS. In this case, the rMOG may be necessary for numerical processing but may not impart any behavioral benefit to congenitally blind individuals. A second possibility is that recruitment of additional cortical regions lends advantages to cognitive processing. Finally, it remains possible that the rMOG does not causally contribute to numerical cognitive processes. The possibility that rMOG activity is entirely epiphenomenal seems less likely since we find that rMOG possesses a relatively fine-grained code for different numerosities. Furthermore, there is evidence that “visual” cortex activity is behaviorally relevant to some higher-cognitive tasks in blindness (Amedi, Floel, Knecht, Zohary, & Cohen, 2004; Merabet et al., 2004). In future work it will be important to test whether the rMOG is functionally relevant to numerical performance in blindness using techniques such as transcranial magnetic stimulation (TMS). Irrespective of the functional relevance of the rMOG to number tasks, the current findings suggest that visual experience expands the cortical territory responsive to numerical information in blindness into the “visual” cortex.

In summary, our results suggest that representations of approximate number are established in the IPS irrespective of sensory experience. However, blindness enhances IPS sensitivity to numerical information presented in auditory numerical sequences. Furthermore, in the absence of vision, neural populations within deaffrented “visual” cortex are capable of developing a numerosity code.

## Acknowledgments

We thank our blind and sighted participants who contributed their time to this research and the F. M. Kirby Research Center for Functional Brain Imaging at the Kennedy Krieger Institute, Rashi Pant and Rita Loiotile for their assistance with data collection. Research reported in this publication was supported by the National Eye Institute of the National Institutes of Health under award number R01EY027352, the Johns Hopkins University Catalyst Award (M.B.) and the National Science Foundation Graduate Research Fellowship DGE-1232825 (S.K.). The content is solely the responsibility of the authors and does not necessarily represent the official views of the National Institutes of Health or National Science Foundation.

## Supplementary Information

**Supplementary Table 1.**
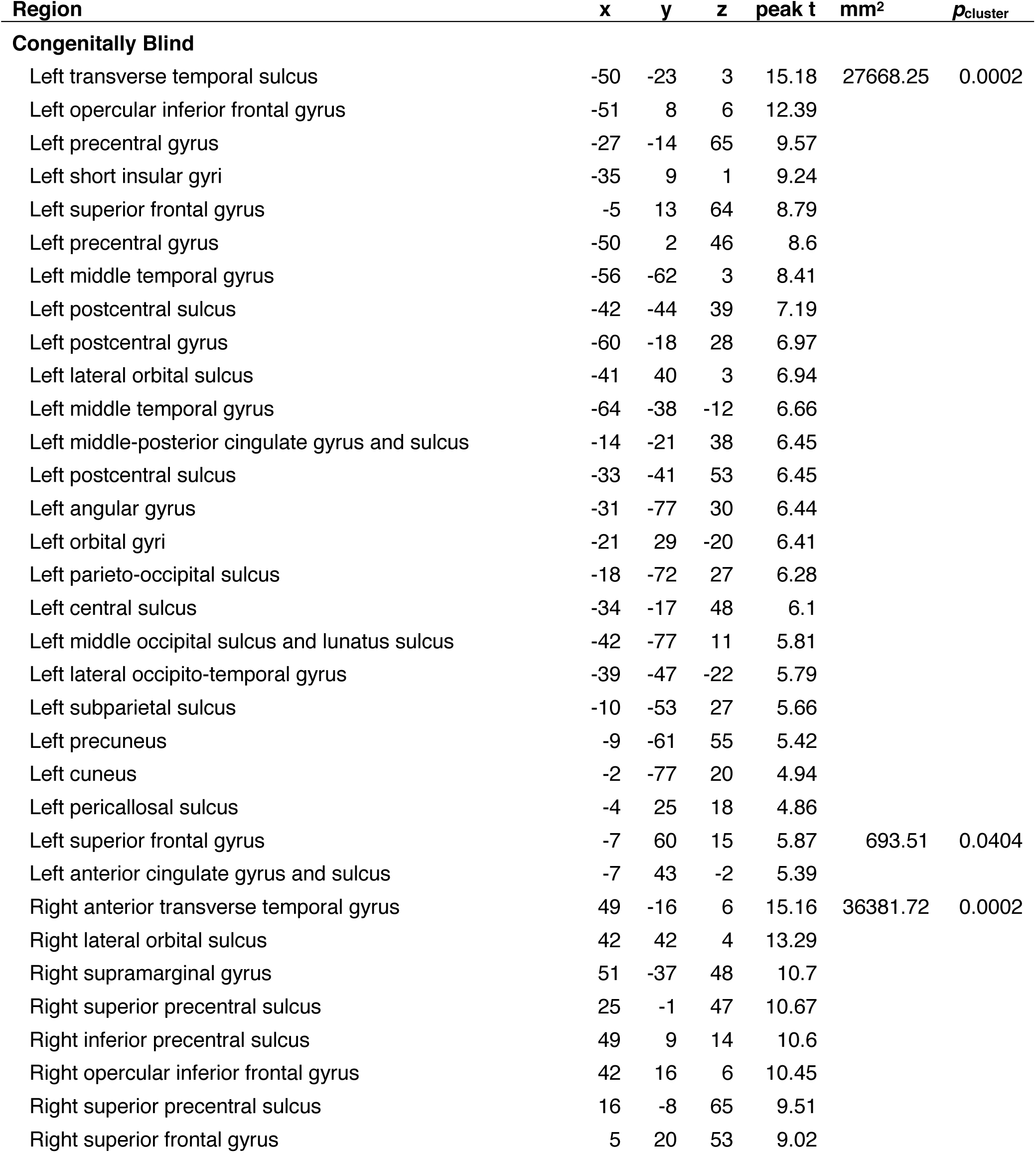
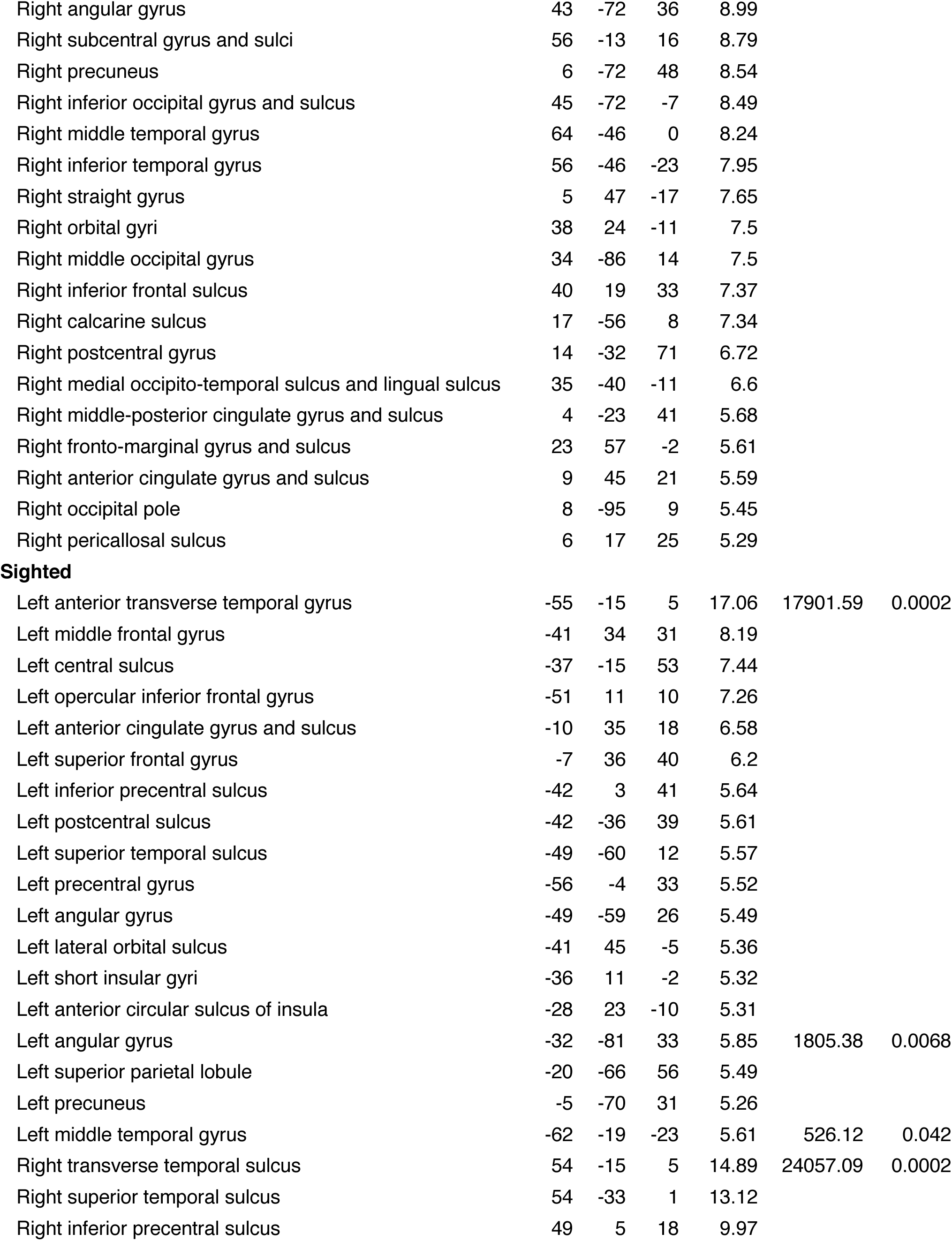
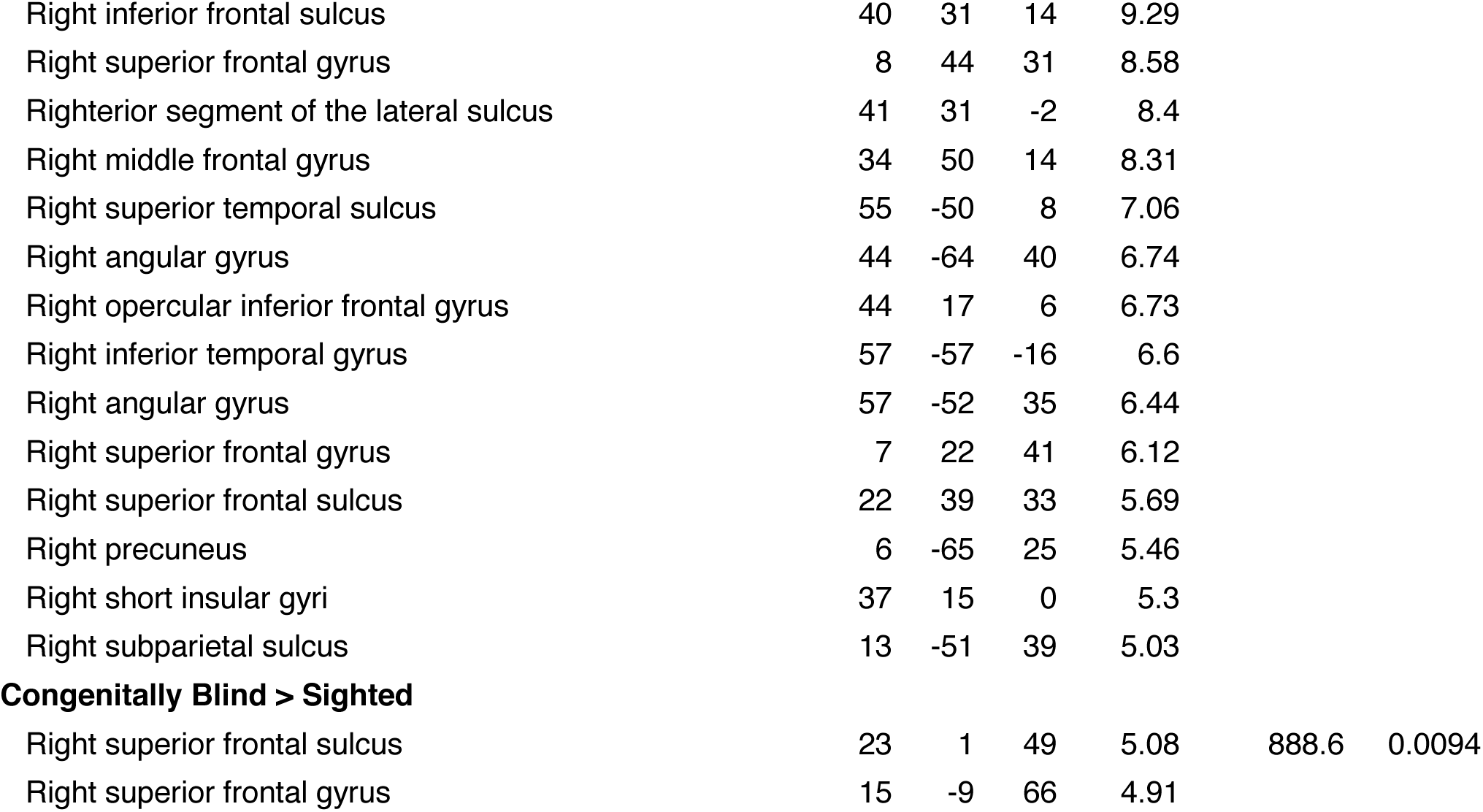
Brain regions in which numerosities could be discriminated.

